# MDM2 stabilization of Notch intracellular domain upon DNA damage plays a major role in non-small cell lung carcinoma response to platinum chemotherapy

**DOI:** 10.1101/2024.04.29.591624

**Authors:** Sara Bernardo, Quentin-Dominique Thomas, Maicol Mancini, Alba Santos, Sylvia Fenosoa Rasamizafy, Amina-Milissa Maacha, Anais Giry, Emilie Bousquet-Mur, Laura Papon, Marion Goussard, Christophe Fremin, Andrea Pasquier, María Rodríguez, Camille Travert, Jean-Louis Pujol, Laetitia K Linares, Lisa Heron-Milhavet, Alexandre Djiane, Irene Ferrer, Luis Paz-Ares, Xavier Quantin, Luis M Montuenga, Hélène Tourriere, Antonio Maraver

## Abstract

Despite major advances in lung cancer clinical management, majority of patients suffering non-small cell lung carcinoma (NSCLC) are treated in first line with platinum in combination with immune checkpoint inhibitors. Although platinum compounds normally display an initial therapeutic effect, relapse constitutes a major challenge in the clinical management of NSCLC patients. Therefore, it is fundamental to understand the relapse underlying mechanisms to find new therapeutic opportunities to improve patients’ survival. Here, we found that different DNA damage inducers increase the protein levels of Notch Intracellular Domain (NICD), i.e., the active form of NOTCH1. Mechanistically, we unveiled that upon platinum treatment, there was a concomitant increase of MDM2 together with NICD, and we also observed an MDM2-mediated ubiquitination and stabilization of NICD. Of note, using patient-derived xenografts displaying intrinsic carboplatin resistance, we demonstrated that the combination of carboplatin with MDM2 and NICD inhibitors increased survival and reduced tumor growth compared with carboplatin in monotherapy. Moreover, in patients with NSCLC who received platinum chemotherapy, MDM2 expression level in the tumor was correlated with poor progression-free survival, further validating MDM2 key role in the response to platinum compounds. Our findings open a therapeutic opportunity for NSCLC patients, the main lung cancer subtype of the leading cause of death by cancer worldwide.

## INTRODUCTION

Lung cancer is the most lethal cancer worldwide with 1.8 million deaths per year (Organization, 2021). The main subtype is non-small cell lung carcinoma (NSCLC) that accounts for around 85% of all lung cancer cases. The three major genomic alterations in NSCLC are oncogenic mutations in the *KRAS* and *EGFR* genes and *ALK* fusions (Skoulidis & Heymach, 2019). The survival of patients who are eligible for targeted therapy, for instance patients with NSCLC harboring EGFR or ALK genetic alterations, has been greatly improved by the development of specific inhibitors, but invariably, most patients will develop resistance at later stages (Girard, 2022; Lin *et al*, 2021). Of note, conventional platinum-based chemotherapy is the main alternative for these patients after the appearance of resistance (Hendriks *et al*, 2023b). Moreover, patients with KRAS-mutated NSCLC, the main oncogenic alteration in NSCLC (Reck *et al*, 2021), are treated with the combination of platinum-based therapy and immune checkpoint inhibitors as first-line therapy, and the same applies to most patients not eligible for targeted therapy. Therefore, platinum chemotherapy, alone or in combination, is still the standard of care for ~60-70% of patients with metastatic NSCLC (Gandhi *et al*, 2018; Paz-Ares *et al*, 2018). However, some patients will develop resistance and other patients will never respond due to intrinsic resistance. Hence, understanding the basis of the response/absence of response to platinum is a major clinical question that needs to be addressed.

Several resistance mechanisms to platinum salts have been proposed, such as decreased cellular drug import, increased cellular detoxification, or enhanced DNA damage repair to name only a few (Huang *et al*, 2021). Regarding the last one, platinum salts form inter- and intra-strand adducts in the DNA double helix that eventually led to double-strand breaks (DSBs), if not properly repaired. DNA DSBs are the most deleterious lesions because they generate genomic instability and trigger the DNA Damage Response (DDR) (Aguilera & Gomez-Gonzalez, 2008). The main DDR sensors are the checkpoint protein kinases ATR and ATM that initiate a signaling cascade to activate cellular programs with the aim of maintaining genome integrity. Depending on the DNA damage extent, the DDR might promote cell cycle arrest to allow DNA repair or might induce cell apoptosis and/or senescence if DNA damage is extreme (Krenning *et al*, 2019).

Activation of the Notch signaling pathway also has been implicated in platinum treatment resistance in different cancers, including lung cancer where carboplatin induces Notch activity through a largely unknown mechanism (Liu *et al*, 2013). Shedding light into this phenomenon could help to identify new ways to target the Notch pathway specifically in NSCLC cells, particularly during platinum treatment. NOTCH is a transmembrane receptor activated by the interaction with transmembrane ligands present on the membrane of neighboring cells. There are four different Notch receptors (NOTCH1-4) and five ligands (JAGGED 1 and 2, and Delta-like 1, 3 and 4) in humans. Ligand binding to the extracellular domain of NOTCH induces a multistep proteolytic cleavage of the receptor. The last step is promoted by the γ-secretase complex leading to the release of the active form of the receptor called NOTCH Intracellular Domain (NICD). After nuclear translocation, in the canonical Notch pathway NICD binds to the transcription factor RBPJ, allowing the displacement of corepressors and the recruitment of coactivators, such as MAML and p300. These factors activate a transcription program that includes among other, HES/HEY family members, Deltex-1 or MYC, depending on the cell type and context (Bray, 2016). We previously demonstrated that the Notch pathway plays a major role in KRAS-driven and in EGFR-driven NSCLC biology as well as resistance to EGFR-targeted therapy (Bousquet Mur *et al*, 2020; Maraver *et al*, 2012). As Notch activity is increased in lung cancer cells upon platinum treatment (Liu *et al*., 2013), we hypothesized that Notch might play a major role also in the resistance to such treatment.

In the present study, we confirmed that the Notch pathway is activated upon platinum-based therapy *in vitro* and expanded also to the *in vivo* setting. Moreover, we found that the DDR, via ATM, promoted such activation. Specifically, we found that MDM2 levels increased upon carboplatin treatment and promoted NICD stability through non degradative ubiquitination. We also demonstrated *in vivo*, using NSCLC patient-derived xenografts (PDX) displaying intrinsic resistance to carboplatin, that the concomitant inhibition of MDM2 and the Notch pathway increases carboplatin effectiveness. Lastly, we also observed that in NSCLC patients treated with platinum chemotherapy, MDM2 levels negatively correlate with progression-free survival (PFS). Altogether, our findings describe a new MDM2-NICD axis induced by carboplatin treatment and open a new therapeutic opportunity for patients with NSCLC.

## RESULTS

### DNA damage induces NICD stabilization

The Notch pathway is upregulated in response to platinum treatment in lung cancer cells (Liu *et al*., 2013), but the underlying molecular mechanisms are largely unknown. Therefore, first we tested whether the Notch pathway was activated only by platinum (carboplatin) or also by other DNA damage inducers currently used for cancer treatment (in particular irinotecan, a topoisomerase II inhibitor, and γ irradiation). As the tumor suppressor p53 plays a major role in the DDR (Abuetabh *et al*, 2022), we used two human KRAS-driven NSCLC cell lines: A549 (harboring wild type p53) and H358 (harboring a homozygous deletion of p53). We incubated A549 and H358 cells with the three genotoxic agents and monitored DDR and Notch pathway activation by measuring the expression levels of phosphorylated (active) ATM and NICD, respectively. All three treatments led to DDR and Notch pathway activation (Figure 1A and Supplemental Figure 1A). This effect was p53-independent because NICD level was similarly increased in both H358 and A549 cells (Figure 1A and Supplemental Figure 1A).

**Figure 1.**
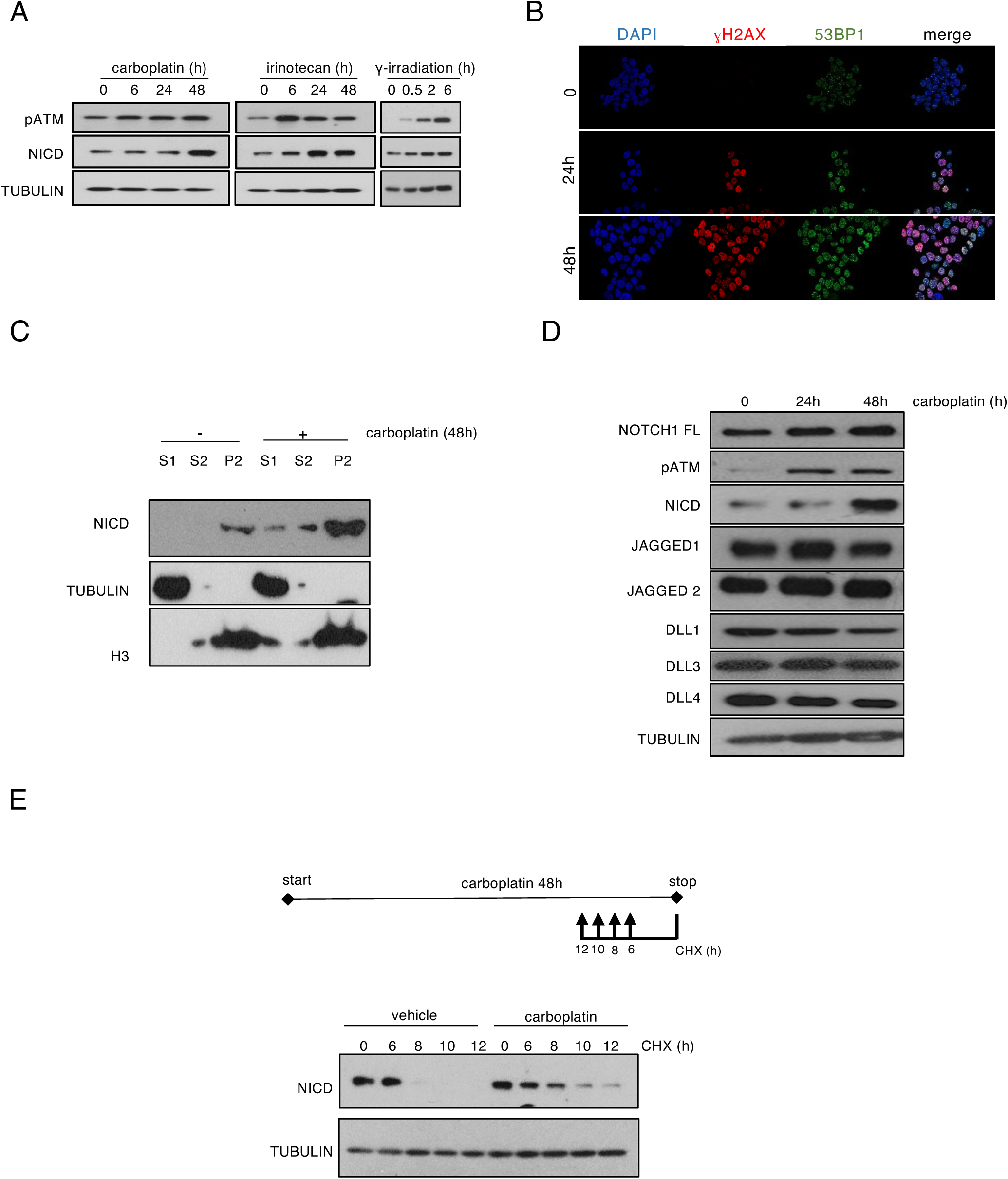
DNA damage induces NICD stabilization. **(A)** Western blotting of the indicated proteins in H358 cells incubated with 100 μM carboplatin, 25 μM irinotecan, or γ-irradiation (4 Gy) for the indicated times. pATM, phosphorylated (activated) ATM. **(B)** Immunofluorescence analysis of γH2AX (in red) and 53BP1 (in green) expression in H358 cells after 24 or 48 hours of incubation with 100 μM carboplatin. Nuclei were stained with DAPI (blue). **(C)** Western blotting of the indicated proteins after cellular fractionation of H358 cells incubated (+) or not (-) with 100 μM carboplatin for 48 hours. S1, cytoplasmic fraction; S2, nucleoplasmic fraction; P2, chromatin fraction; TUBULIN, loading control of S1; H3, loading control for P2. **(D)** Western blotting of the indicated proteins in H358 cells incubated with 100 μM carboplatin for the indicated times. **(E)** Western blotting of the indicated proteins in H358 cells incubated or not with 100 μM carboplatin for 48 hours and with 50 μg/μl cycloheximide (CHX) for the indicated times before the end of carboplatin treatment. Upper panel: Schematic representation of the experiment design.

We then focused on carboplatin; the molecule clinically relevant for NSCLC treatment. We first confirmed that DNA DSBs were produced upon incubation with carboplatin, as indicated by the formation of γH2AX and 53BP1 foci (Figure 1B and Supplemental Figure 1B). Following γ secretase-mediated cleavage of the NOTCH transmembrane domain, NICD is translocated to the nucleus to exert its transcriptional program (Bray, 2016). Therefore, to further confirm the Notch pathway activation upon carboplatin treatment in our cellular models, we used subcellular fractionation and analyzed NICD levels on different subcellular compartments. NICD strongly accumulated in the chromatin fraction of platinum-treated H358 and A549 cells (thus p53-independent), validating the increased Notch activity upon DNA damage in our system (Figure 1C and Supplemental Figure 1C). Taken together, our data demonstrate that in H358 and A549 cells, carboplatin induced both DDR and the Notch pathway. Therefore, this experimental system can be used to understand how the Notch pathway is increased upon platinum treatment.

NICD levels can be increased through enhanced NOTCH1 processing on the membrane promoted by increased ligand and/or full-length NOTCH1 expression, or by NICD stabilization (Bray & Gomez-Lamarca, 2018). To determine by which mechanism NICD increased its levels upon DNA damage induction, we analyzed the protein expression of all Notch ligands (i.e., JAGGED1, JAGGED2, DLL1, DLL3 and DLL4) and of NOTCH1 full length upon carboplatin incubation. In both H358 and A549 cells, phosphorylated ATM and NICD were consistently induced upon carboplatin treatment, but this was not the case for full-length NOTCH1 or its ligands (Figure 1D and Sup Figure 1D), favoring an increase in NICD stability to explain its increase in protein levels. To determine whether NICD half-life was increased upon DNA damage, we incubated both cell lines with carboplatin or vehicle for 48 h and added cycloheximide (a translation inhibitor) for different amounts of time before the end of the exposure to carboplatin (Figure 1E). Importantly, NICD stability was increased in cells incubated with carboplatin compared to vehicle and this was true in both H358 and A549 cells (Figure 1E and Sup Figure 1E), thus independently of p53 expression.

Taken together all our results so far indicated that DNA damage increased NICD stability.

### NICD stabilization upon DNA damage is ATM-dependent

We showed above that three different genotoxic agents increased NICD levels, and hence, we hypothesized that the DDR was responsible for NICD increased stability upon carboplatin treatment. Two main DNA damage sensor kinases orchestrate the DDR, i.e., ATM and ATR, hence we used specific inhibitors to test their possible role in NICD stabilization upon DNA damage. Thus, we pre-incubated H358 and A549 cells with KU55933 (ATM inhibitor) and ETP-46464 (ATR inhibitor) (Thanasoula *et al*, 2012) for 2 hours before carboplatin addition and both kinases were efficiently inhibited as indicated by the decreased levels of phosphorylated ATM and ATR respectivelly (Figure 2A-B). ATR inhibition did not decrease NICD accumulation upon DNA damage, but conversely, ATM inhibition drastically reduced NICD levels in H358 and A549 cells incubated with carboplatin (Figure 2A-B). To confirm that ATM controlled NICD stability upon carboplatin treatment, we performed cycloheximide pulse-chase experiments in the presence of carboplatin and ATM inhibitor. In accordance with our previous experiments, carboplatin stabilized NICD levels for 8 hours (Figure 2C-D). Conversely, ATM inhibition dramatically reduced NICD half-life and it was barely detectable after 4h (Figure 2C-D).

**Figure 2.**
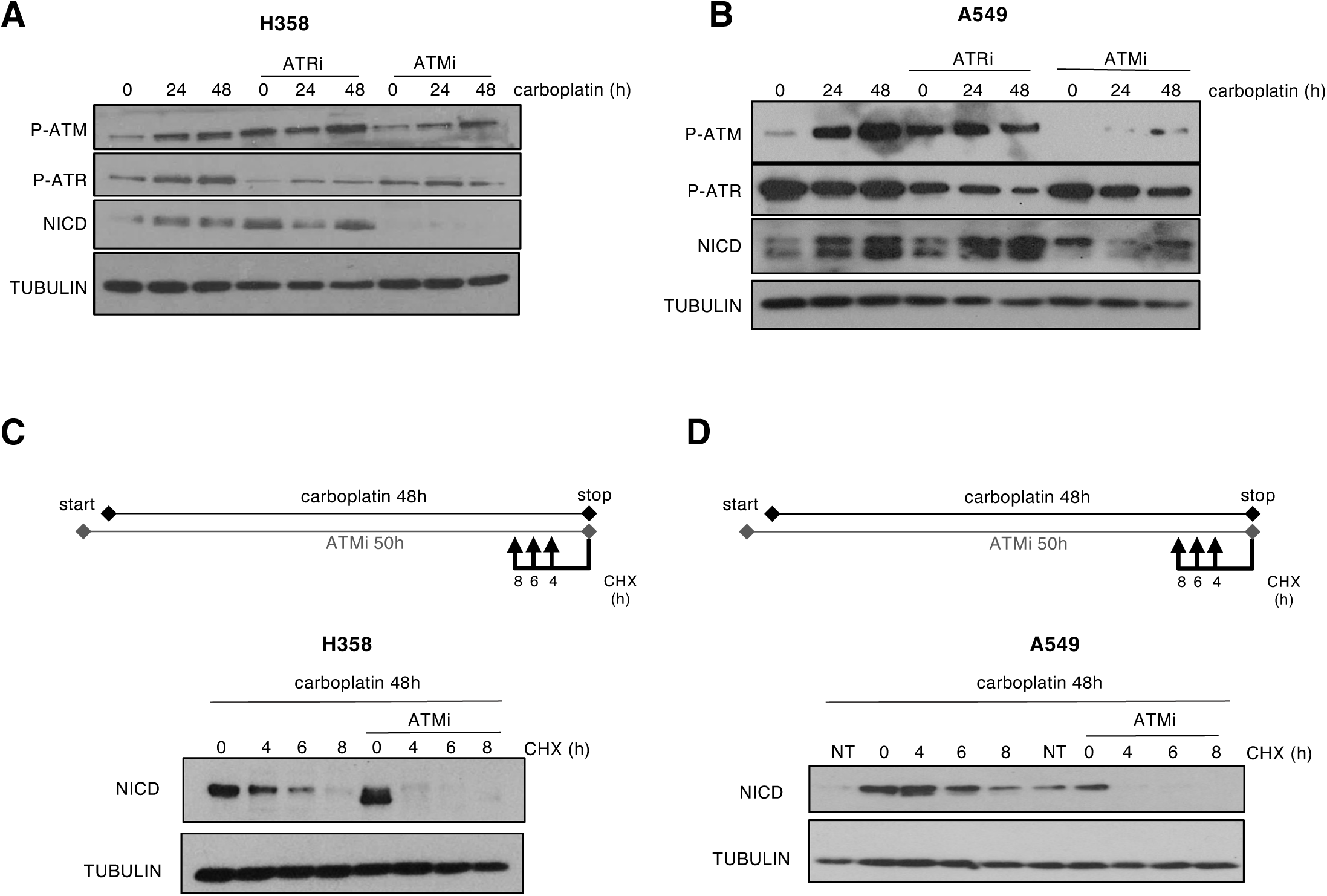
NICD stabilization upon DNA damage is ATM-dependent. **(A-B)** Cells were incubated with 100 μM carboplatin and with/without 10 μM KU-55933 (ATM inhibitor) or 5 μM ETP-64646 (ATR inhibitor) for the indicated time followed by immunoblotting to detect the expression of the indicated proteins in H358 **(A)** and A549 **(B)** cells. P, phosphorylated. **(C-D)** H358 **(C)** and A549 **(D)** cells were incubated with 100 μM carboplatin for 48 hours and with or without 10 μM KU-55933 (ATM inhibitor). 50 μg/μl cycloheximide was added for the indicated time before the end of carboplatin treatment. Upper panel: Schematic representation of the experimental design.

Our findings suggest that ATM is required for increasing NICD stability upon DNA damage.

### NICD is ubiquitinylated upon DNA damage and MDM2 promotes its stabilization

To finely control Notch signaling, NICD is regulated by various post-translational modifications, including ubiquitination that promotes NICD degradation (Lai, 2002). Therefore, to identify if DNA damage induces NICD ubiquitination, we used a well-characterized system in which ectopic expression of His6-tagged ubiquitin allows the purification of *de novo* ubiquitinated proteins (Treier *et al*, 1994). We generated a NOTCH1 delta extracellular domain construct (NOTCH1-deltaE) (see Materials and Methods for details) to express a ligand-independent but secretase-dependent NOTCH1 protein. This allowed increasing NICD expression in the cells, but not in a supraphysiological manner as it is frequently the case upon direct NICD overexpression. We co-transfected both plasmids in 293T cells and then incubated them with carboplatin as before. Due to the increased protein levels of NICD upon DNA damage (Figure 1A and Supplemental Figure 1A), we were expecting a lower ubiquitination, but surprisingly, it was strongly increased (Figure 3A and Supplemental Figure 3A). Interestingly, previous work described that E3 ligase MDM2 interacted with and ubiquitinated NICD, leading to hyperactivation of the Notch pathway in the absence of any type of DNA damage (Pettersson *et al*, 2013). Even more MDM2 is an important DDR downstream actor and displays p53-independent functions (Bouska & Eischen, 2009), making it a good E3 ligase candidate for DNA damage-induced NICD ubiquitination. Thus, we first analyzed MDM2 protein levels in H358 and A549 cells upon carboplatin exposure. Intriguingly, MDM2 expression increased upon DNA damage and decreased upon ATM inhibition in both cell lines (i.e., p53-independent), mirroring both phosphorylated ATM and NICD patterns (Figure 3B and Supplementary Figure 3B). Our new data suggest that MDM2 could be responsible for NICD increased stability upon DNA damage and therefore, we analyzed the effect of MDM2 loss of function in this setting.

**Figure 3.**
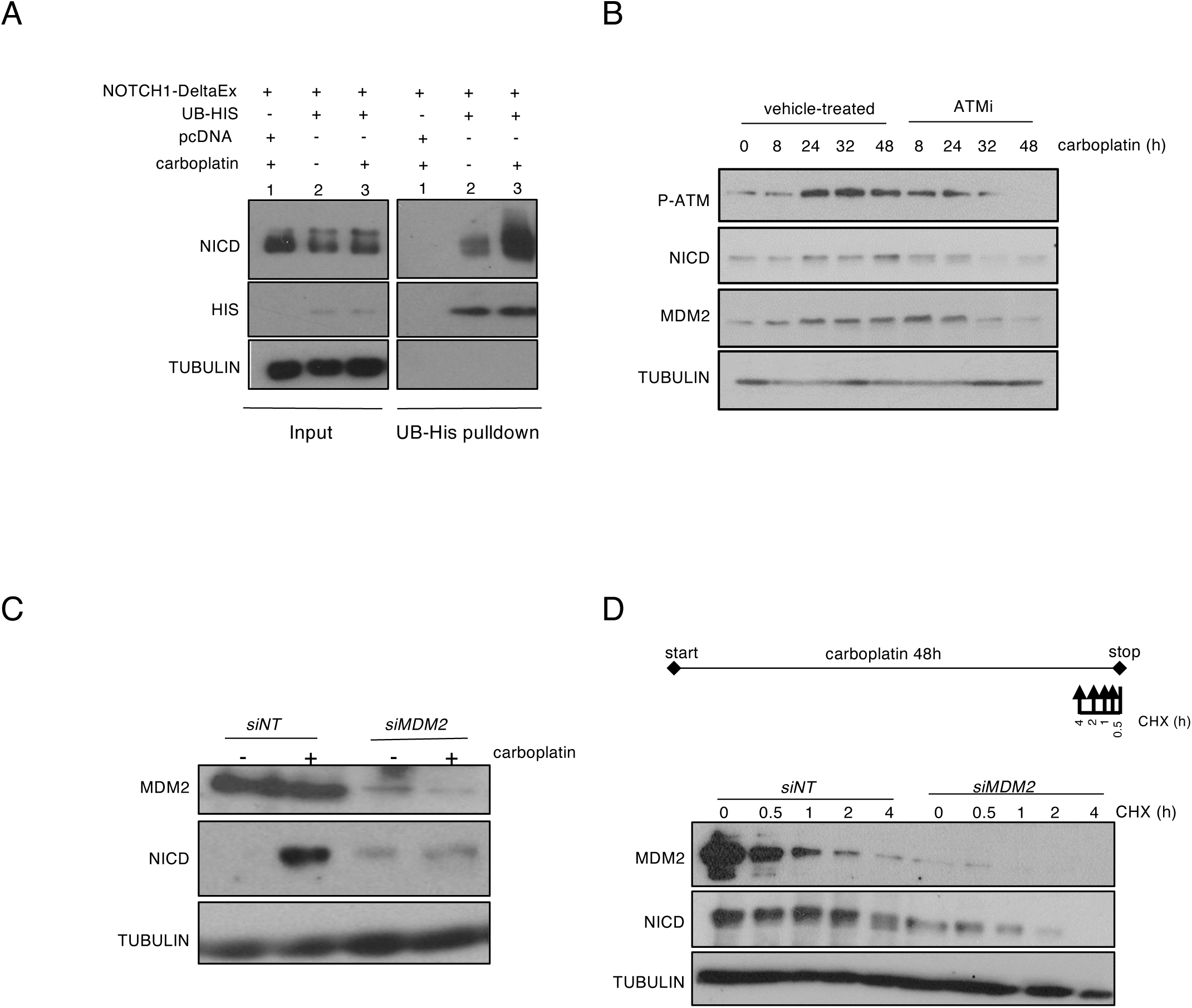
NICD is ubiquitinylated upon DNA damage and its stabilization is MDM2-dependent. **(A)** 293T cells were transfected to express NOTCH1-DeltaEx, 6His-Ubiquitin (UB-HIS) or empty vector (pcDNA), as indicated, and incubated (+) or not (-) with carboplatin for 48h. Histidine pull-down was performed and NICD ubiquitination was analyzed by western blotting. INPUT, total cell extracts. **(B)** Western blotting of the indicated proteins in H358 cells incubated with 100 μM carboplatin with or without 10 μM KU-55933 (ATM inhibitor) for the indicated time. **(C)** Western blotting of the indicated proteins in H358 cells in which *MDM2* was silenced with a *siRNA* against MDM2 (*siMDM2*) or not (non-targeting *siRNA*; *siNT*). At 6 h post-transfection, 100 μM carboplatin was added to the cells for 48 h. **(D)** At 6 h post-transfection, H358 cells transfected with *siMDM2* or *siNT* were incubated with 100 μM carboplatin for 48 h and with 50 μg/μl of cycloheximide for the indicated time before the end of carboplatin treatment. Upper panel: schematic representation of the experimental design.

In accordance with our previous data, in control cells (*siNT-*treated), NICD protein levels increased upon carboplatin treatment. Conversely, and validating our hypothesis, in *siMDM2-*treated cells, NICD was not increased upon carboplatin addition in either H358 nor A549 cells (Figure 3C and Supplementary Figure 3C, respectively). To further demonstrate that MDM2 promoted NICD stability upon carboplatin treatment, we monitored NICD half-life in *siMDM2-* and *siNT*-treated cells incubated with cycloheximide and carboplatin. In both H358 and A549 cells subjected to MDM2 loss of function, NICD stabilization was strongly decreased after DNA damage (Figure 3D and Supplementary Figure 3D).

Taken together, our data shows that upon DNA damage, NICD is ubiquitinated and MDM2 is required to increase its stability.

### The MDM2 E3-ligase activity is essential for ubiquitination and increased stability of NICD

Then, to determine whether MDM2 was sufficient to stabilize NICD, we ectopically expressed wild type MDM2 together with NOTCH1-deltaE in human 293T cells. Even more, to demonstrate that its E3 ligase activity was required for such stabilization, we also expressed an E3 ligase-dead MDM2 mutant (C464A point mutation in the RING domain) (Riscal *et al*, 2016). Of note, in the absence of any DNA damage, ectopic expression of wild type MDM2, but not of the ligase-dead MDM2 mutant, was sufficient to increase NICD levels in 293T cells, also in the chromatin fraction (Figure 4A).

**Figure 4.**
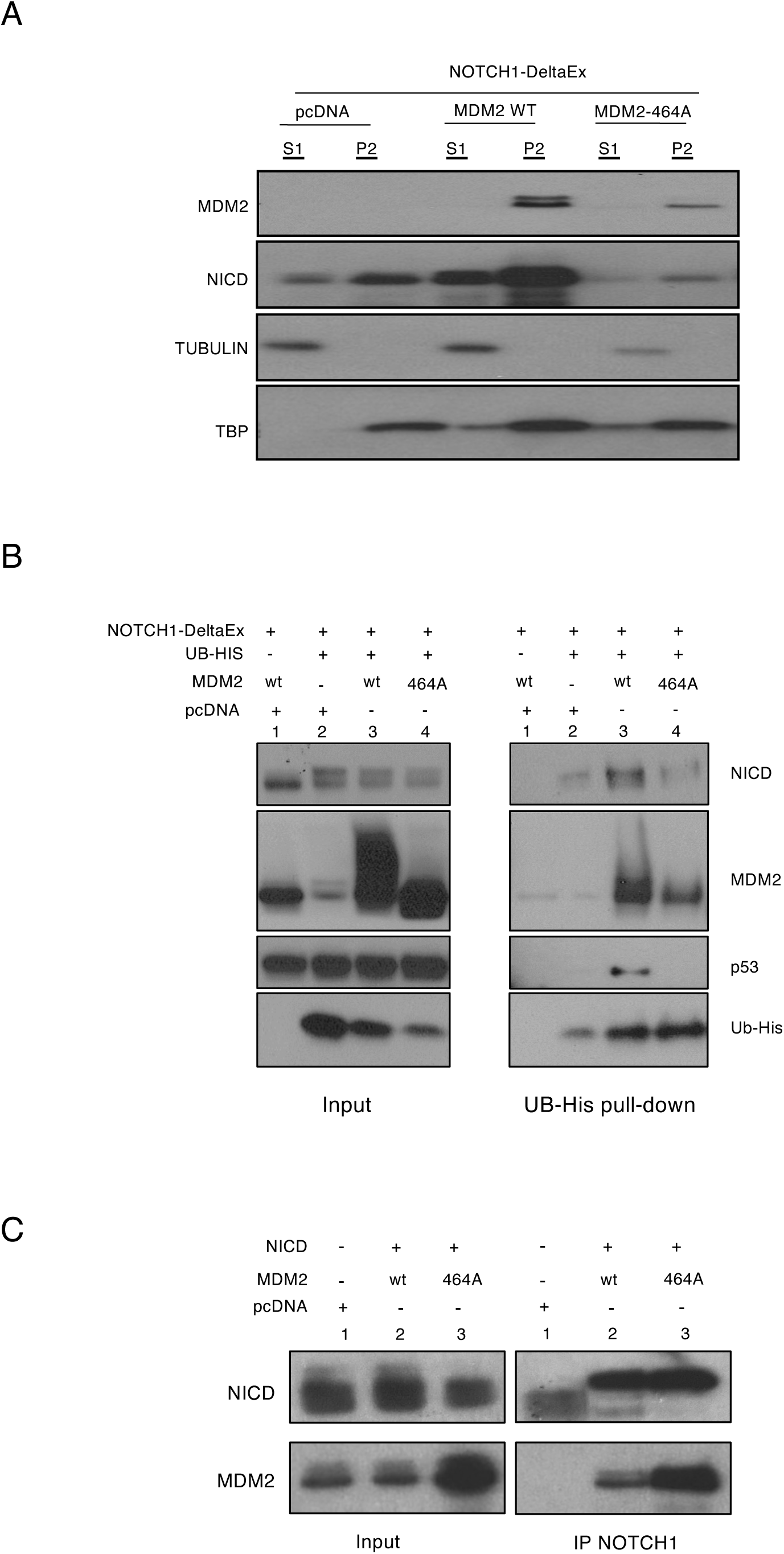
MDM2 E3-ligase activity is required for NICD ubiquitination. **(A)** 293T cells were transfected with NOTCH1-DeltaEx and then co-transfected with empty vector (pcDNA), wild type (WT) MDM2 or MDM2 464A mutant for 48h. Then, NICD levels were analyzed by western blotting after cell fractionation. S1, cytoplasmic fraction; P2, chromatin fraction. TUBULIN, loading control for the S1 fraction; TBP: loading control for the P2 fraction. **(B)** 293T cells were transfected with NOTCH1-DeltaEx and then co-transfected with 6xtag-His-Ubiquitin (UB-HIS), empty vector (pcDNA), MDM2 WT or MDM2 464A mutant for 48h followed by histidine affinity pull-down and western blotting. INPUT: total cell extracts. (**C**) 293T cells were transfected with NOTCH1-DeltaEx, empty vector (pcDNA), MDM2 WT, or MDM2 464A mutant for 48h. Cell extracts were used for NOTCH1 immunoprecipitation and the levels of the indicated proteins were measured by western blotting. INPUT: total cell extracts.

To formally prove that NICD was a MDM2 E3 ligase substrate, we ectopically expressed NOTCH1-deltaE, His6-tagged ubiquitin, and wild type or ligase-dead MDM2 in 293T cells. In the absence of ectopic MDM2, NICD was weakly ubiquitinated (Figure 4B, line 2 on the right panel and supplemental Figure 4, lines 2 and 3 on the right panel). Importantly, NICD ubiquitination was strongly increased in cells that expressed ectopic wild type MDM2 (Figure 4B, line 3 on the right panel and supplemental Figure 4, lines 4 and 5 on the right panel), but not in cells that expressed the ligase-dead MDM2 C464A mutant (Figure 4B, line 4 on right panel and supplemental Figure 4, lines 6 and 7 on the right panel). To control the experiment, we showed that ubiquitination of p53 (the best-known target of MDM2) was increased by ectopic expression of wild type MDM2 but not by expression of the MDM2 C464A mutant (Figure 4B, third line on right panel).

To assess whether a MDM2 and NICD interaction shown previously (Pettersson *et al*., 2013) occurs in our setting, but also whether the E3 ligase function was implicated in this interaction, we performed co-immunoprecipitation experiments using 293T cells that ectopically expressed NOTCH1-deltaE and either wild type or MDM2 C464A mutant. MDM2 and NICD were similarly co-precipitated independently of MDM2 mutational status (Figure 4C). Our data confirmed that MDM2 and NICD do interact and also showed that although the E3 ligase activity was crucial for increasing NICD ubiquitination upon DNA damage, it was not required for the MDM2-NICD interaction.

Altogether, our findings indicate that the MDM2-NICD axis is induced upon DNA damage.

### Combined inhibition of NICD and MDM2 enhances platinum effectiveness and increases survival *in vivo*

As we established that Notch pathway is activated via MDM2 upon carboplatin treatment, we investigated whether carboplatin effect could be enhanced by inhibiting the MDM2-NICD axis. To increase the clinical relevance of our approach, we used a NSCLC patient-derived xenograft (PDX) that displayed partial intrinsic resistance to carboplatin (Ferrer *et al*, 2018). We used SP141, an MDM2 inhibitor that induces its degradation by forcing its autoubiquitination without affecting p53 interaction (Wang *et al*, 2014), and dibenzazepine (DBZ), a Notch inhibitor that we previously demonstrated to be effective in NSCLC PDXs *in vivo* (Bousquet Mur *et al*., 2020).

After the PDX subcutaneous graft in nude mice, we let the tumors grow to 200 mm^3^ before randomization into three treatment groups: vehicle, carboplatin, and carboplatin with the MDM2 and Notch inhibitors. We administered carboplatin on Wednesday, and SP141 and DBZ from Monday to Friday (two cycles). To confirm proper target inhibition, we first used a small group of mice (n = 4 mice/group). At the end of the second treatment cycle, we assessed protein levels in PDX-derived tumors by immunohistochemistry (IHC) using the histoscore (H-score) (Hirsch *et al*, 2003) and found that: 1) γH2AX H-score was increased in both carboplatin treatment groups (compared with vehicle), confirming that carboplatin at the dose used induced DNA damage in the tumors *in vivo*; 2) in accordance with our *in vitro* findings, MDM2 expression was also increased upon carboplatin treatment compared with vehicle, but decreased when carboplatin was combined with inhibition of the MDM2-NICD axis; 3) to monitor Notch activity we used a known read-out of this pathway, i.e., HES1 and found that its H-score was not increased by carboplatin alone but was drastically decreased (as expected) in the platinum plus MDM2-NICD axis inhibition group. The HES1 result in the carboplatin group was intriguing because as mentioned above, in the absence of carboplatin, HES1 is a suitable read-out by IHC of Notch pathway activation in KRAS- and EGFR-driven NSCLC (Bousquet Mur *et al*., 2020; Maraver *et al*., 2012). Hence, we concluded that HES1 is not a reliable read-out of Notch activity increase promoted by DNA damage, but it is still a good marker of Notch inhibition; 4) Ki67 (cell division marker) H-score was increased in the carboplatin group (compared with vehicle) and decreased in the combination group compared with carboplatin (and not different from vehicle); 5) cleaved caspase 3 expression (C3A) was increased in the combination group compared to carboplatin-treated group, and further increased (by 2.4 X-fold) when compared to vehicle-treated mice, suggesting that MDM2-NICD axis inhibition could potentiate carboplatin effect in this model (Figure 5A). As HES1 was not a reliable read-out of Notch activity in carboplatin treated tumors, and NICD expression cannot be detected by IHC, we determined NICD expression in tumors by western blotting and found that in accordance with our in vitro data, it was strongly increased in the carboplatin group, but not in the combination one (Figure 5B). To the best of our knowledge, this is the first demonstration of Notch activation *in vivo* upon DNA damage.

**Figure 5.**
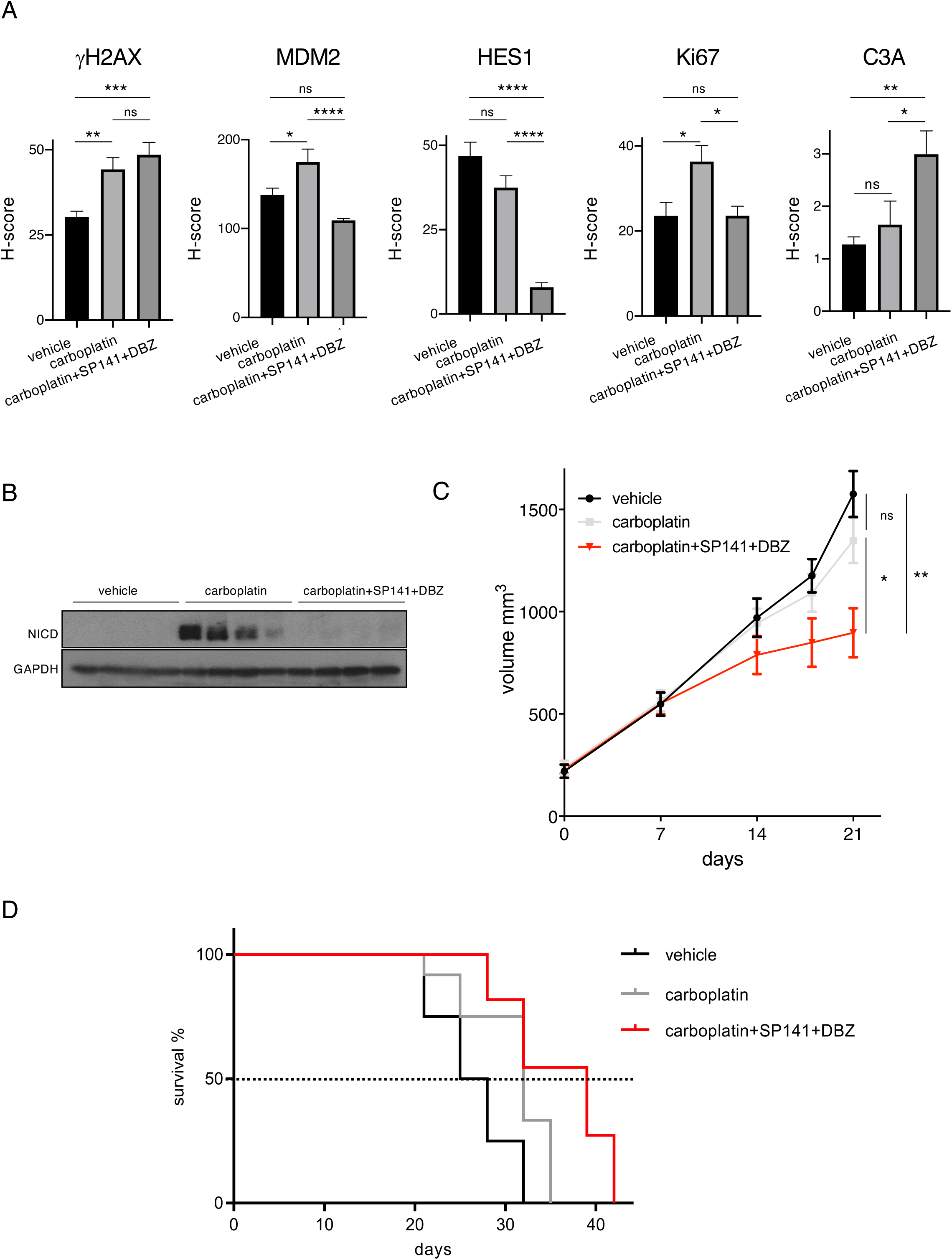
Combined inhibition of NICD and MDM2 enhances platinum effectiveness and increases survival *in vivo*. **(A)** IHC analysis of γH2AX, MDM2, HES1, Ki67 and cleaved caspase 3 (C3A) in PDX (TP57) cell xenografts from nude mice treated with vehicle (Methocel, n = 16 measures from 4 different tumors), carboplatin alone (n = 16 measures from 4 different tumors), or the DBZ (Notch inhibitor) + SP141 (MDM2 inhibitor) + carboplatin combination (n = 16 measures from 4 different tumors). (**B**) The levels of the indicated proteins were measured by western blotting in the same mouse groups as in A (n = 4 different tumors). (**C**) Growth of PDX TP57 tumors on nude mice treated with vehicle (Methocel, n = 12), carboplatin (n = 12) or the combination of carboplatin + SP141 + DBZ (n = 12). The *y* axis shows the tumor volume measured with a caliper. (**D**) Survival analysis in the same animals described in C. The *y* axis shows the percentage of surviving animals and the *x* axis the days after treatment. Statistical significance was determined with the Log rank (Mantel-Cox) test. Vehicle vs carboplatin, *p* = 0.08; vehicle vs carboplatin + SP141 + DBZ, *p* = 0.0005; carboplatin vs carboplatin + SP141 + DBZ, *p* = 0.02.

Then, we repeated the experiment using a larger number of mice, i.e., 12 mice per arm, but using the same treatment groups and schedule as in Figure 5A. We measured tumor growth in all mice until the first one reached the end point (tumor volume of 1200 mm^3^) that was day 21 for the vehicle- and carboplatin-treated groups. Tumor growth was much lower in the combination group (carboplatin plus inhibition of the MDM2-NICD axis inhibition) compared with the carboplatin and vehicle groups, while carboplatin alone effect on tumor growth was mild and not significant compared to vehicle (Figure 5C). We also analyzed for survival and we showed that carboplatin had a small but significant effect compared with vehicle (Figure 5D). Remarkably, survival was strongly increased in the carboplatin plus MDM2-NICD inhibition compared with either vehicle or with carboplatin monotherapy (Figure 5D). In particular, the median survival was 26.5, 32 and 39 days for the vehicle-, carboplatin- and carboplatin plus MDM2-NICD-treated groups, respectively.

Taken all together, our preclinical data establish that MDM2-NICD axis inhibition enhances carboplatin therapeutic effect in NSCLC *in vivo*.

### MDM2 expression correlates with poor progression-free survival in patients with NSCLC treated with platinum compounds

To further confirm the clinical relevance of the MDM2-NICD axis implication in NSCLC, we analyzed primary tumor samples from a cohort of 41 patients with NSCLC following platinum administration, alone or in combination with other treatments (see Material and Methods for details). As we showed that HES1 was an unreliable readout of the Notch pathway activation in the presence of DNA damage, we only analyzed for MDM2 expression (using H-score, as in the PDX model). We ranked patients based on MDM2 expression on tumor cells, from the highest to the lowest H-score, and separated them into four quartiles. We analyzed progression free survival (PFS) in the group with the lowest MDM2 expression, (i.e., the lowest quartile) and the group with the highest expression (i.e., the highest quartile). The median PFS were 29.5 months in the lowest MDM2 expression group and 5.4 months in the highest MDM2 expression group (hazard ratio 5.734, 95% confidence interval, 1.505–21.841, *p* = 0.004) (Figure 6A). This robust differences in PFS suggests that MDM2 plays a major role in the response to platinum-based chemotherapy in patients with NSCLC.

**Figure 6.**
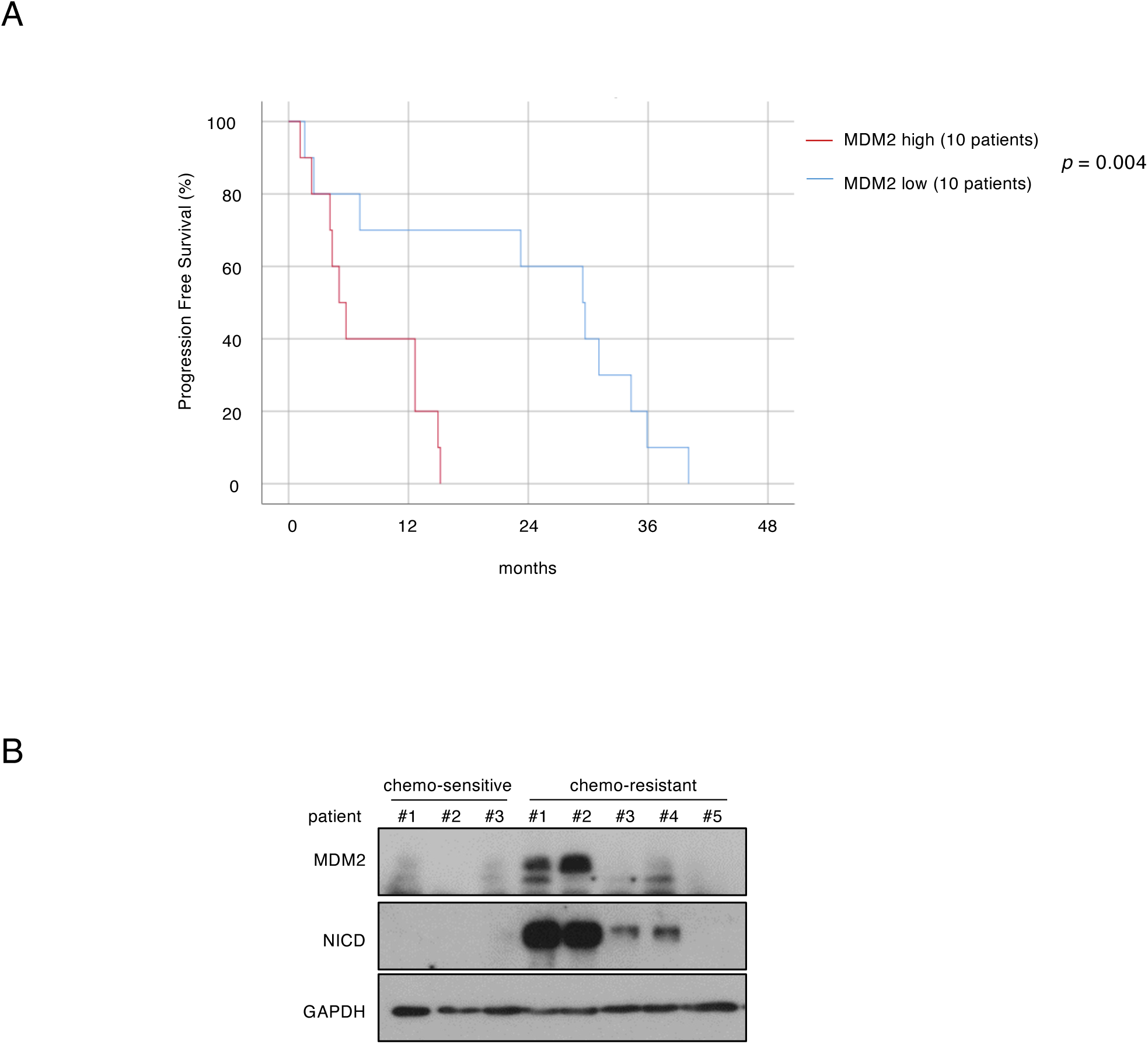
MDM2 expression correlates with poor progression-free survival in patients with NSCLC treated with platinum-based chemotherapy. **(A)** Progression-free survival of patients with NSCLC treated with platinum drugs (n = 41) in function of MDM2 expression in the tumor assessed by IHC: highest quartile (n = 10) versus lowest quartile (n = 10) of MDM2 expression. Statistical significance was determined with the Log rank (Mantel-Cox) test, *p* = 0.004. **(B)** Immunoblotting of the indicated proteins in tumor samples from three chemo-sensitive and five chemo-resistant patients with NSCLC treated with platinum drugs (neoadjuvant chemotherapy).

Lastly, we assessed NICD protein levels in a smaller cohort of patients with NSCLC and available frozen tumor tissue after surgery. All these patients received platinum-based neoadjuvant chemotherapy before surgery and were classified as chemo-sensitive (n = 3) and chemo-resistant (n = 5) in function of the tumor cell clearance in adjacent lymph nodes. In four of the five chemo-resistant patients, MDM2 and NICD expression levels were high, and interestingly, the expression levels were similar between both proteins, indicating that the MDM2 control of NICD stability we observed in vitro, may also happen in NSCLC patients. Conversely, MDM2 and NICD were not detectable in the three chemo-sensitive patients (Figure 6B). These data strongly suggest that the MDM2-NICD axis plays an important role in the response to platinum in patients with NSCLC.

Our clinical data confirmed and expanded the mechanistic findings showing that the MDM2-NICD axis plays a major role in the DDR and carboplatin effectiveness in NSCLC.

## DISCUSSION

In the last years, massive research efforts have been made to find new therapeutic option for NSCLC, the deadliest cancer worldwide (Organization, 2021). These efforts unveiled unprecedent and innovative treatments, such as immune checkpoint and KRAS^G12C^ inhibitors. Nevertheless, platinum-based chemotherapy remains the most used treatment for patients with early to late-stage lung cancer (Hendriks *et al*, 2023a; Hendriks *et al*., 2023b), and overcoming resistance against this therapeutic option is an unmet medical need. Therefore, we must increase our knowledge on the changes occurring in lung cancer cells upon treatment with platinum-based drugs in order to develop new drug combinations to tackle the appearance of resistance.

In the current study, we first discovered that DNA damage activates the Notch pathway. We then showed that ATM, one of the main downstream sensors of DDR, is implicated in Notch signaling activation. As previous studies showed that Notch inhibits ATM (Vermezovic *et al*, 2015), we hypothesize that under physiological conditions, the ATM-NOTCH crosstalk may establish a negative feedback loop to regulate ATM activation upon DDR. Then, we showed that upon incubation with DNA damage-inducing agents, NICD ubiquitination increased concomitant with its stability. This was a counterintuitive observation, as it is normally accepted that ubiquitination promotes NICD degradation (Lai, 2002). This important result led us to MDM2 because, to the best of our knowledge, it is so far the only E3 ubiquitin ligase reported to interact with and activate NICD (Pettersson *et al*., 2013). Importantly, our study highlights that the MDM2-mediated stabilization of NCID is p53-independent, although it is widely known that MDM2 also binds to p53 promoting its ubiquitination and degradation to tightly control its activity. This is in agreement with recent studies showing that some MDM2 roles in cancer are p53-independent (Arena *et al*, 2018; Cisse *et al*, 2020; Riscal *et al*., 2016) and also with *in vivo* studies demonstrating that cisplatin effect in lung cancer cells is p53-independent (Oliver *et al*, 2010).

One key observation in our work is that MDM2 accumulated upon DNA damage induction in an ATM-mediated manner. Previous studies showed that upon DDR, ATM phosphorylates MDM2 at S394 to promote p53 activation (Gannon *et al*, 2012), whereas AKT phosphorylation of MDM2 at S166 and S186 increases its translocation to the nucleus (Mayo & Donner, 2001), as observed in our study. As ATM is required for AKT activation upon exposure to ionizing radiation (Viniegra *et al*, 2005), we hypothesize that the ATM-induced AKT-mediated effect on MDM2 is more prevalent than the direct effect of ATM on MDM2. Our findings also showed that upon DNA damage, MDM2 interacts with and ubiquitinates NICD, unveiling a previously undescribed signaling axis that operates upon carboplatin treatment. We also verified that MDM2 stabilizes NICD in the absence of DNA damage and it could explain, at least in part, why MDM2 amplification is an independent factor of poor prognosis in patients with NSCLC (Dworakowska *et al*, 2004). Moreover, by analyzing NSCLC samples from a subset of patients treated with platinum-based chemotherapy we found that NICD and MDM2 expression levels were positively correlated and that the MDM2-NICD axis was activated only in non-responder patients. Lastly, in another cohort of patients with NSCLC and treated with platinum-based chemotherapy we found that MDM2 expression level was inversely correlated with PFS. Therefore, our data strongly suggest that high MDM2 expression, by amplification or other mechanisms, will affect the response to platinum-based chemotherapy in NSCLC by stabilizing NICD.

Our preclinical data in mice harboring NSCLC PDXs with carboplatin intrinsic resistance, further support this hypothesis because the combination of carboplatin and MDM2-NICD axis inhibition increased survival and limited tumor growth compared to carboplatin alone. This indicates that targeting the MDM2-NICD axis is a good strategy to boost carboplatin effect in NSCLC.

Another interesting finding was that upon incubation with carboplatin to induce DNA damage, HES1 was not a reliable biomarker of Notch activation, unlike in untreated EGFR- (Bousquet Mur *et al*., 2020) and KRAS-driven NSCLC (Maraver *et al*., 2012). Additional work is required to understand the reason behind this unusual observation, but we anticipate that it might be related to the uncoupling of HES1 and NICD signaling upon DNA damage. For instance, HES1 is present in the Fanconi anemia core complex but not NICD (Tremblay *et al*, 2008), and so, we could hypothesize that other pathways regulate HES1 expression upon DNA damage.

Overall, our data suggest a new therapeutic opportunity for patients who do not respond or become resistant to platinum-based chemotherapy, the most common treatment for the leading cause of death by cancer worldwide, i.e., lung cancer (Organization, 2021).

## MATERIAL AND METHODS

### Cell culture and proliferation assay

The H358 (KRAS^G12C^ mutation, homozygous deletion of p53), A549 (KRAS^G12S^, mutation, wild type p53) and 293T cell lines were cultured in RPMI 1640 medium containing 10% fetal calf serum, 10% antibiotics in humidified atmosphere of 5% CO_2_ at 37°C and were tested for absence of mycoplasma contamination. Cycloheximide (Sigma) was added at 50μg/ml at the indicated time points. Carboplatin (ACCORD Healthcare, France) was used at 100 μM for 48h. KU55933 (ATM inhibitor (Selleckchem) and ETP-46464 (ATR inhibitor) were used at 10 μM and at 5 μM, respectively, and added 2 hours before carboplatin treatment. Irinotecan (Fresenius Kabi) was used at 25 μM for 48 hours. For γ-irradiation, cells were exposed to a single dose of 4 grays (Gy), at a dose rate of 2.7 Gy/min using a Xstrahl XenX preclinical irradiator.

### Protein extraction and western blotting

Cell extracts were prepared by incubating cells on ice in 50 mM Tris–HCl buffer (pH 8.0), supplemented with 150 mM NaCl, 5 mM EDTA, 0.5% DOCNa, 1% SDS, 1% TritonX-100, phosphatase and protease inhibitor cocktail (Sigma, USA, #P5726 and #P8340) for 30 min. Protein concentrations were measured with the Pierce TM BCA Protein Assay Kit (ThermoFisher Scientific, USA). Proteins were separated by SDS-PAGE, transferred to PVDF membranes and analyzed with the following antibodies against: NICD (#4147, 1:500), phosphorylated ATM (#13050,1:1000), phosphorylated ATR (#2853, 1:1000), JAG1 (70109S, 1 :1000), JAG2 (2210T, 1:1000), DLL3 (78110S 1:1000), DLL4 (96406S, 1:1000), full-length NOTCH1 (3608S, 1:1000), GAPDH (2118S, 1:5000), H3 59715S, 1:5000), all from Cell Signaling Technology, Inc, and also DLL1 (QC16739-41705, Antiva Biosciences, 1:1000), MDM2 (#4B2C1.11 and #4B11 Merck Millipore, 1:2000), tubulin (#T9026, Sigma, 1:10000), poly-histidine (H1029 Sigma, 1:1000), p53 (sc-126 Santa Cruz Technology, 1:1000), and TBP (TFIID sc-56794, Santa Cruz Technology, 1:500). Secondary antibodies were horseradish peroxidase-linked anti-rabbit (#7077, Cell Signaling Technology, 1:10000 dilution), or anti-mouse (#7076, Cell Signaling Technology, 1:10000 dilution) IgG. Antibody binding was detected by chemiluminescence using the ECL detection system (GE Healthcare) or ECL Plus (for NICD) (GE Healthcare).

### Generation of NOTCH1-DeltaEx and cell transfection

The NOTCH1-DeltaEx plasmid was constructed by adding the NOTCH transmembrane domain into a NICD plasmid, kindly provided by Dr Bijan Sobhian (Yatim *et al*, 2012). The whole sequence was cloned into the Gateway entry vector pDONR221 using specific primers containing the AgeI and MluI restriction sites combined with attB specific sites. Using the Gateway system (Invitrogen), the NOTCH1-DeltaEx construct was then cloned into the Gateway destination vector pCLX-R4-DEST-R2, a kind gift from Dr. Patrick Salmon (Addgene plasmid # 45956).

Wild type MDM2 and MDM2^C464A^ were previously described (Riscal *et al*., 2016) and also the His6-tagged ubiquitin construct (Treier *et al*., 1994). The three constructs were provided by Dr Laetitia Linares, author of this study.

For MDM2 loss of function, the siGENOME SMARTpool against MDM2 (L-003279-00-0005) was obtained from Thermo Scientific Dharmacon RNAi Technologies and transfected in H358 and A549 cells according to the manufacturer’s instructions. After 6h, cells were incubated with carboplatin for 48h.

### Immunofluorescence

Cells were grown on 12 mm glass coverslips and treated as described. Cells were then washed in PBS, fixed in 2% formaldehyde at room temperature for 15 min and permeabilized in 0.5% Triton X-100 in PBS for 10 min. Cells were incubated in blocking buffer for 1 h, followed by primary antibodies against γH2AX (the same antibody used for wester blotting) and 53BP1 (1:1000, MAB3802 Merck). Cells were washed and incubated with secondary anti-rabbit-Alexa Fluor 488 (1:500, Thermo Fisher Scientific) and anti-mouse-Cy3 (1:500, Thermo Fisher Scientific) antibodies at room temperature for 1 h. After washing, cells were incubated with DAPI (1 μg/ml) and mounted onto glass slides using DAKO Fluorescent Mounting Medium (Agilent Technologies S3023). Images were acquired using an Upright ZEISS AXIO Imager.M2 with Apotome 2.

### Chromatin fractionation

Chromatin fractionation was performed as previously described (Smits *et al*, 2006). Approximately 3×10^6^ cells were washed in PBS and resuspended in 200 μl of solution A (10 mM HEPES pH 7.9, 10 mM KCl, 1.5 mM MgCl_2_, 0.34 M sucrose, 10% glycerol, 1 mM DTT, protease and phosphatase inhibitors). Triton X-100 was added to a final concentration of 0.1%, cells were incubated on ice for 5 min, and the cytoplasmic (S1) and nuclear fractions (P1) were harvested by centrifugation at 1300 g for 4 min. Isolated nuclei were washed in solution A, lysed in 150 μl of solution B (3 mM EDTA, 0.2 mM EGTA, 1 mM DTT, protease and phosphatase inhibitors), and incubated on ice for 10 min. The soluble nuclear (S2) and chromatin fractions were harvested by centrifugation at 1700g for 4 min. Isolated chromatin (P2) was then washed in solution B, spun down at 10000 g, and resuspended in 150 μl 1X Laemmli buffer.

### Ubiquitination analysis

293 T cells were transfected with the indicated plasmids and recovered by trypsinization; 10% of all harvested cells was used to prepare the total cell extract for western blot analysis, as described above. The remaining cells were resuspended in 10ml guanidine denaturing buffer (6M guanidine chloride, 20mM Tris, 0.01M imidazole, 0.5mM DTT, 0.1% iodoacetamide, 0.5% Triton X-100) and incubated at 4°C under agitation for 1h. Extracts were incubated with 50μl of Ni-NTA Agarose beads/sample (HisPur™ Ni-NTA Resin, ThermoScientific) at 4°C for 2h under agitation to promote the binding of histidine with the nickel beads. Then, the bead/protein complexes were gravity-filtered on a sintered column (Fisher Scientific, Cytiva 17-0435-01), washed in 5ml guanidine denaturing buffer followed by washing first with 20mM imidazole buffer (2X PBS, 20mM imidazole, 0.25% Triton X-100) and then with 50mM imidazole buffer (2X PBS, 50mM imidazole, 0.25% Triton X-100). Beads were recovered in 50μl of 4X Laemmli buffer, denatured at 95°C for 10 min and subjected to immunoblotting.

### Immunoprecipitation

Extracts were prepared as described in the Chromatin fractionation method, but only the isolated chromatin was used, resuspended in micrococcal nuclease buffer, incubated with micrococcal nuclease (M0247S, Biolabs) at 37° for 10 min, and resuspended in the same volume of lysis buffer (20mM Tris pH 7.5, 150mM NaCl, 5% glycerol, 0.5% NP40, 0.2mM EDTA). For IP, extracts were diluted three times in dilution buffer (20mM Tris pH 7.5, 100mM NaCl, 0.2mM EDTA) and incubated with 5μl of anti-NOTCH1 antibody (Ab27526, Abcam) at 4°C, with agitation, overnight. This was followed by incubation with 50 μl of Dynabeads™ Protein G magnetic beads (Thermo Fisher Scientific) for 1h to bind the immunoprecipitated complexes to the beads. After several washes in wash buffer (20mM Tris pH 7.5, 150mM NaCl, 0.25% NP40, 0.2mM EDTA), the bead-antibody-protein complexes were resuspended in 50 μl of 4X Laemmli buffer and denatured at 95° for 10 min. The supernatants containing the immunoprecipitated proteins were separated on SDS-PAGE gels and analyzed by immunoblotting.

### Mice

The PDX was generated in Luis Paz-Ares’ laboratory (one of the study authors) at the Instituto de Biomedicina de Sevilla (IBIS). The PDX was from a NSCLC that harbored KRAS (G12A) and p53 (P151R) mutations and displayed intrinsic resistance to carboplatin (Ferrer *et al*., 2018). A piece of 0.5 mm^3^ was implanted into the right flank of 6-week-old athymic Nude-Foxn1 female mice. Drug treatments were started when tumors reached 200 mm^3^. Mice were killed when tumors reached 1200 mm^3^. Dibenzazepine (DBZ) was obtained from Syncom (#SIC-020042, The Netherlands). It was administered at 2.2 mg/kg/day 5 days per week (Monday–Friday) by intraperitoneal (ip) injection. Carboplatin (10mg/ml, ACCORD Healthcare, France) was administered at 50 mg/kg/day 1 day per week (Wednesday) by ip, as already described (Oliver *et al*., 2010). SP141 was synthesized by the SynBio3 platform (Gilles Subra’s team, Montpellier) and was administered at 30 mg/kg/day 5 days per week by ip (Cisse *et al*., 2020; Wang *et al*., 2014). Tumor growth was monitored using a caliper.

Animal procedures were performed according to protocols approved by the French national committee of animal care.

### Immunohistochemistry

PDX tumors were fixed, embedded in paraffin and stained with hematoxylin and eosin or used for IHC. IHC of HES1, Ki67, cleaved caspase 3, γ-H2AX and MDM2 was performed by the RHEM platform as previously described (Bousquet Mur *et al*., 2020; Cisse *et al*., 2020). For patient samples, immunochemistry was done by ICM Translational Research Unit (N° CORT: ICM-CORT-2022-15) with anti-HES1 (clone D6P2U, #11988, Cell Signaling Technology, 1:100 dilution) and -MDM-2 (clone IF2, # MABE340, Millipore, 1:500 dilution) antibodies.

For each tumor, four 10X magnification fields were scored using the Qpath software, providing a total of 16 measures per tumor. Protein expression was evaluated according to the staining intensity and percentage of positive cells using the H-score (Hirsch *et al*., 2003). Staining intensity was scored as negative (0), weak (1), moderate (2), or strong (3) and the percentage of positive cells was reported for each staining intensity. The H-score ranged from 0 to 300 and was calculated by taking into account the percentage of positive tumor cells and the staining intensity (Hirsch *et al*., 2003).

### Patients and ethical considerations

For the PFS analysis (Figure 6A), surgical primary lung cancer samples were obtained from the University Clinic of Navarra (CUN), Spain. The cohort included 41 patients who received the cancer diagnosis between 2001 and 2021 and who met the following inclusion criteria: diagnosis of NSCLC, surgical resection of the primary lung tumor, availability of clinical data and chemotherapy treatments that included platinum alone or in combination. Exclusion criteria: presence of another primary tumor in the 5 years before surgery, excluding non-melanoma skin tumor.

The protocol for obtaining samples was approved by the CUN Ethics Committee and all patients signed a written informed consent. PFS was defined as the period between the diagnosis and the first relapse. The Kaplan-Meier method was used for the survival analysis and the log rank test to compare groups. The Cox proportional hazards model was used to determine the hazard ratios (HR) with the IBM SPSS Statistics 25 software.

For the western blot analysis (Figure 6B), primary NSCLC tumors were obtained from Hôpital Arnaud de Villeneuve, Montpellier, France. All patients had NSCLC and had received at least three cycles of platinum-based chemotherapy. Patients were classified as responders or non-responders, based on the absence and/or persistence of the initial lymph node metastases, respectively. This study was approved by the Committee for the Protection of Persons of each institution and by the French National Agency for Medicines and Health Products Safety (ANSM).

All patients signed an informed consent to use their tumor samples for research.

### Statistical analysis

Unless otherwise specified, the data are presented as means ± S.E.M. One way analysis of variance (ANOVA) followed by Tukey’s post hoc test was performed to assess the significance of expression levels in IHC. for changes in size of tumors a repeated measures two-way ANOVA followed by Tukey’s post hoc test was performed. Kaplan-Meier survival curves were analyzed with the log rank test.

* *p* ≤ 0.05; ** *p* ≤ 0.01; *** *p* ≤ 0.001, **** p ≤ 0.0001

## ACKNOWLEDGMENTS

We thank Eric Chevet and David Santamaría for helpful discussion and critical reading of the manuscript. Elisabetta Andermarcher professionally edited the manuscript. We thank the IRCM animal facility members for their outstanding work. We acknowledge the “Réseau d’Histologie Expérimentale de Montpellier” for processing our animal tissues, histology techniques and expertise. We acknowledge the ICM/IRCM experimental irradiation facility supported by SIRIC, granted by “INCa-DGOS-INSERM-ITMO Cancer_18004” for the irradiator. S.B. was supported by a French Ministry of Education and Research fellowship. Work in A.M.’s laboratory is supported by the Institut National du Cancer (INCa_9257 and INCa_11554) and by SIRIC Montpellier (INCa_Inserm_DGOS_12553). This project received also support from Ligue Nationale contre le cancer (Comité de l’hérault, R20028FF) and from INCa-Cancéropôle GSO (R20056FF). This work was also supported by FIMA, CIBERONC (CB16/12/00443), Spanish Ministry of Economy and Innovation and Fondo de Investigación Sanitaria Fondo Europeo de Desarrollo Regional (PI22/00451 to L.M.M.). The funders had no role in the study design, data collection and analysis, decision to publish, or preparation of the manuscript.

