## Supplemental Material for "MDM2 stabilization of Notch intracellular domain upon DNA damage plays a major role in non-small cell lung carcinoma response to platinum chemotherapy"

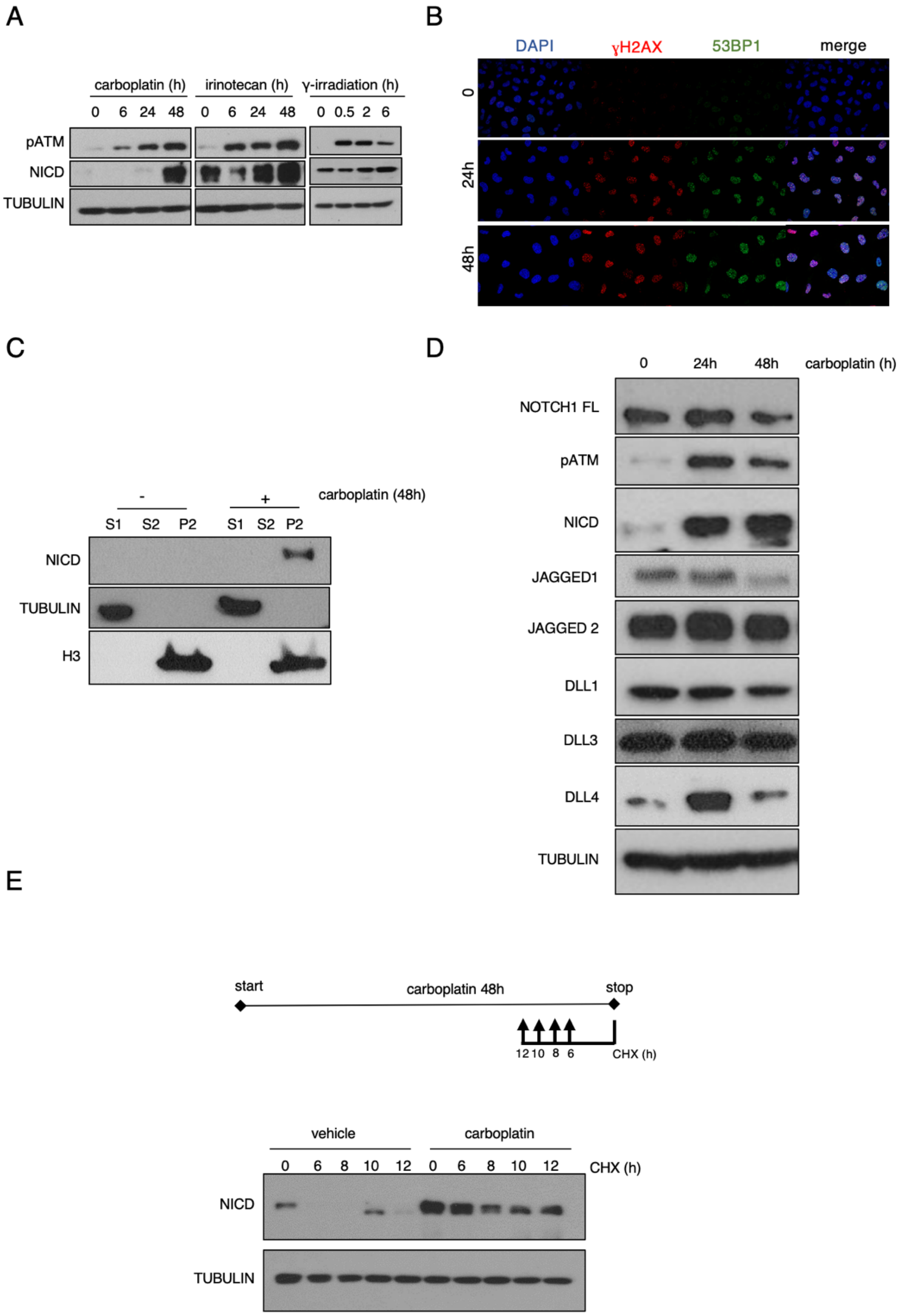


**Supplemental Figure 1. DNA damage induces NICD stabilization.**

**(A)** Western blotting of the indicated proteins in A549 cells incubated with 100 μM carboplatin, 25 μM irinotecan, or γ-irradiation (4Gy) for the indicated time. pATM, phosphorylated (activated) ATM. **(B)** Immunofluorescence analysis of γH2AX (in red) and 53BP1 (in green) in A549 cells after 24 or 48 hours of incubation with 100 μM carboplatin. Nuclei were stained with DAPI (in blue). **(C)** Western blotting of the indicated proteins after cellular fractionation of A549 cells incubated (+) or not (-) with 100 μM carboplatin for 48 hours. S1, cytoplasmic fraction; S2, nucleoplasmic fraction; P2, chromatin fraction; TUBULIN, loading control for S1; H3, loading control for P2. **(D)** Western blotting of the indicated proteins in A549 cells incubated with 100 μM carboplatin for the indicated time. **(E)** Western blotting of the indicated proteins in A549 cells incubated or not with 100 μM carboplatin 48 hours and with 50 μg/μl cycloheximide (CHX) for the indicated time before the end of carboplatin treatment. Upper panel: Schematic representation of the experimental design.

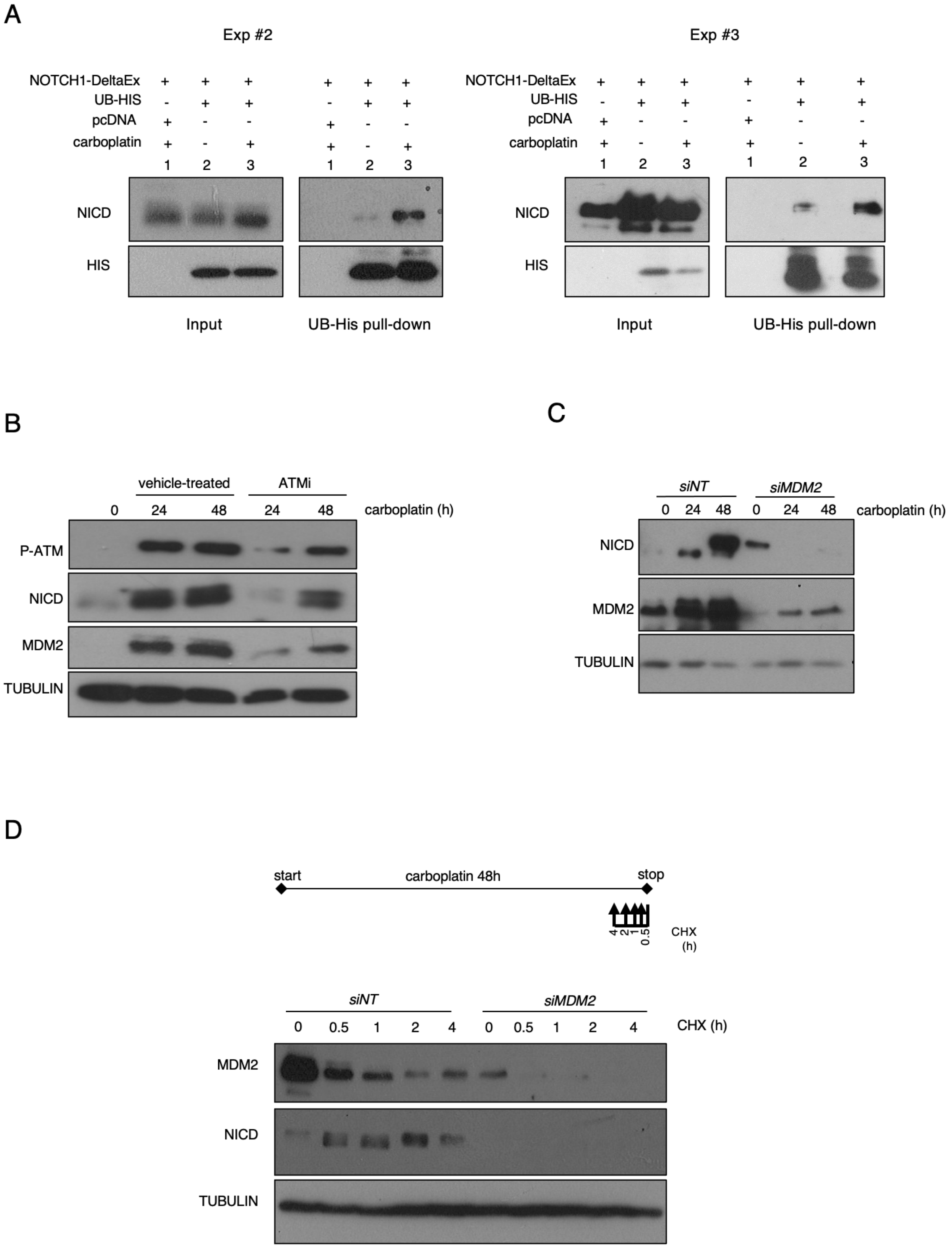


**Supplemental Figure 3. NICD stabilization is MDM2-dependent.**

**(A) Experiments done as in Fig3A (Exp #2 and Exp #3); (B)** Western blotting of the indicated proteins in A549 cells incubated with 100 µM carboplatin for the indicated time, with or without 10 µM KU-55933 (ATM inhibitor). **(C)** Western blotting of the indicated proteins in A549 cells transfected with a non-targeting *siRNA* (*siNT*) or with a *siRNA* against MDM2 (*siMDM2*) for 6 hours, followed by addition of 100 µM carboplatin for the indicated time. **(D)** At 6 h post-transfection, A549 cells transfected with *siMDM2* or *siNT* were incubated with 100 µM carboplatin for 48 h and with 50 µg/µl of cycloheximide for the indicated time before the end of carboplatin treatment. Upper panel: schematic representation of the experimental design.

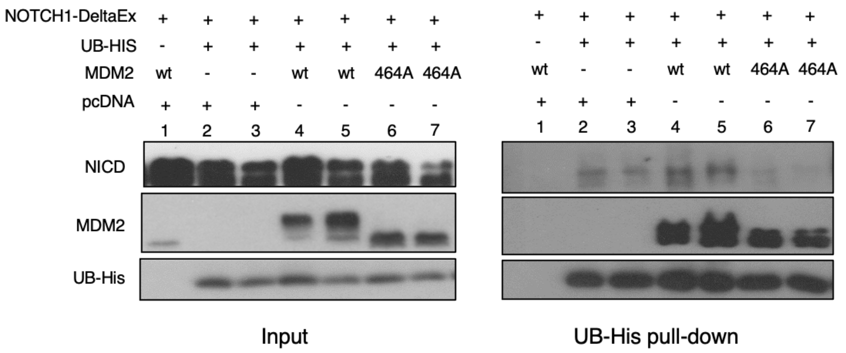


**Supplemental Figure 4. MDM2 E3-ligase activity is required for NICD ubiquitination**.

293T cells were transfected with NOTCH1-DeltaEx and then co-transfected with 6xtag-His-Ubiquitin (UB-HIS), empty vector (pcDNA), MDM2 WT or MDM2 464A mutant for 48h followed by histidine affinity pull-down and western blotting. INPUT: total cell extracts.

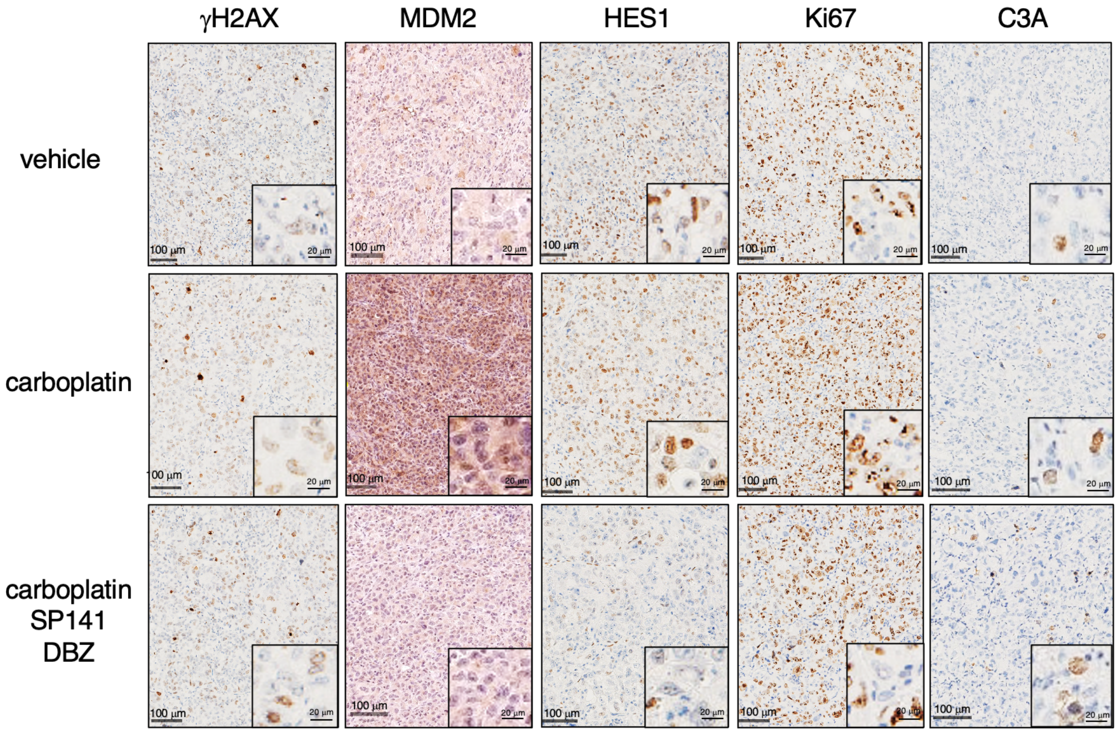


**Supplemental Figure 5. Examples of IHC of the graphs presented in Figure 5A.**
